# Screening and machine-learning assisted prediction of translation-enhancing peptides reducing ribosomal stalling in *Escherichia coli*

**DOI:** 10.1101/2025.07.17.665026

**Authors:** Teruyo Ojima-Kato, Gentaro Yokoyama, Hideo Nakano, Michiaki Hamada, Chie Motono

**Author notes:** The person to whom correspondence. T.O.K and G.Y. contributed equally to this work.

## Abstract

We previously reported that the nascent SKIK peptide enhances translation and alleviates ribosomal stalling caused by arrest peptides (APs) such as SecM and polyproline when positioned immediately upstream of the APs in both *Escherichia coli* in vivo and in vitro translation systems. In this study, we performed a comprehensive screening of translation-enhancing peptides (TEPs) using a randomized artificial tetrapeptide library. The screening was based on the ability of the peptides to suppress SecM AP-induced translational stalling in *E. coli* cells. Various TEPs exhibiting a range of translation-enhancing activities were identified. In vitro translation analysis suggested that the fourth amino acid in the tetrapeptide plays a key role in reducing SecM AP-mediated stalling. Furthermore, we developed a machine learning model using a random forest algorithm to predict TEP activity. The predicted values showed a strong correlation with experimentally measured activities. These findings offer a compact peptide toolkit and a data-driven approach for mitigating AP-induced ribosome stalling, with potential applications in synthetic biology.

## Introduction

Efficient protein synthesis is fundamental to synthetic biology and the importance is also becoming increasing for sustainable bio-research and industry. However, despite advances in gene design and codon optimization, synthesis of protein of interest (POI) can often be influenced by a variety of factors including promoter, nucleotide sequence of the mRNA, and the availability of tRNAs, thereby limiting protein production yields and compromising the functionality of synthetic circuits^1^

Translation, one of the critical steps of protein synthesis, is known to be influenced by the sequence of the nascent polypeptide chain itself^2,3^. Emerging evidence has shown that specific nascent peptide sequences, referred to as arrest peptides (APs), can interact with the ribosomal exit tunnel to induce ribosome stalling during elongation^4–6^. These AP-mediated stalls play pivotal roles in regulatory circuits that fine-tune gene expression in response to environmental or physiological signals^3,7^.

A prominent example is the SecM AP (FSTPVWISQAQGIRAGP), found in *Escherichia coli*, which regulates the translation of secA, an essential component of the Sec protein translocation system^4^. The SecM AP stalls ribosomes in a manner dependent on its specific amino acid sequence, particularly within the arrest motif^8^. In addition, stretches of consecutive proline residues (polyproline motifs) are also known to cause ribosome stalling due to the limited availability of prolyl-tRNA and the slow kinetics of proline incorporation^9^. While these regulatory mechanisms are biologically advantageous in certain contexts, they pose significant challenges in biotechnology and synthetic biology, where efficient and uninterrupted translation is crucial for high-yield protein production. Overcoming ribosome stalling has thus become a focus of research aimed at enhancing the efficiency of recombinant protein expression systems^10,11^.

Our research group previously reported that the insertion of an “SKIK peptide tag” composed of the four amino acids Ser-Lys-Ile-Lys at the N-terminus of difficult-to-express proteins, enhances protein production not only in both *E. coli* in vivo and in vitro systems but also in *Saccharomyces cerevisiae*^12–14^. So far, this peptide tag has been proved to be useful in increasing protein synthesis by enhancing translation while the mechanism remaining unclear^15–17^. More recently we demonstrated that a short nascent peptide sequence, SKIK, when positioned immediately upstream of an AP such as SecM or a polyproline motif, can alleviate ribosome stalling and contribute increased protein production in *E. coli*^18,19^. On the other hand, Kobo et al. reported ribosomal stalling by AP could be attenuated by randomly chosen several tetrapeptides^20^. Herynek et al. reported more direct screening approach to get specific N-terminal peptide to increase soluble production of each POI by using GFP fused POI and fluorescent activated cell sorter^21^.

While the molecular basis of this phenomenon remains largely unclear, these findings suggest that short peptide sequences could be strategically employed as translation enhancing modules in synthetic constructs to improve protein production. However, the sequence features that confer such translation-promoting activity remain poorly understood, and the potential for discovering new translation enhancing peptides (TEPs) has not been systematically explored. In the context of synthetic biology, identifying and utilizing TEPs offers a modular and programmable approach to overcoming translation barriers. By integrating TEPs into genetic constructs, synthetic biologists can fine-tune translational efficiency, optimize metabolic pathway fluxes, and enhance the production of valuable biomolecules such as enzymes, therapeutic proteins, and biomaterials^22,23^.

In recent years, machine-learning techniques have greatly advanced our ability to navigate protein sequence space^24–28^. Bayesian-optimization frameworks, for instance, have been used to accelerate the functional engineering of proteins^29^. Generative AI models now offer a complementary route, enabling de novo sequence generation^30–33^. However, both approaches depend on large training datasets and are therefore poorly suited to the design of very short peptides—such as four-amino-acid sequences—where data are inherently scarce.

In this study, we aimed to comprehensively identify novel TEPs capable of alleviating ribosome stalling caused by the SecM AP in *E. coli.* To achieve this, we constructed an artificial randomized tetrapeptide library fused with SecM AP followed by superfolder green fluorescent protein (sfGFP) gene. Our screening identified a variety of tetrapeptides with translation enhancing activity of varying strengths. Furthermore, we applied machine learning methods, including a random forest algorithm, to predict TEP candidates based on sequence features, offering a data-driven strategy for optimizing synthetic biology designs. Our findings provide valuable insights into the design principles governing translation efficiency in the *E. coli* protein expression platform.

## Results and Discussion

### Screening of TEPs in *E. coli* from the constructed library

Screening scheme of translation enhancing peptides in this study was summarized in Figure 1. To construct the plasmid library, totally 1.4 × 10^5^ *E. coli* HST08 clones of transformants were obtained, and the diversity of the library was confirmed by sequencing the randomized (NNK)_4_ positions of several clones (data not shown). The pET22b-(NNK)_4_-SecM AP-sfGFP plasmids extracted from the pooled *E. coli* clones were then used for transformation of *E. coli* BL21(DE3) for protein expression. Of the total 1.3 × 10^5^ transformants, about 0.1 % showed fluorescence as shown in Figure 1B. Although the total number of screened tetrapeptide sequences did not reach the full 160,000 possible combinations, the scale was sufficient for an in vivo screening system in terms of order of magnitude. Further analysis of total 217 clones including all positive clones with various fluorescence intensities and some clones showing less fluorescence revealed that they corresponded to 157 unique peptide sequences after removing duplicate sequences. Strength of sfGFP was different depending on the peptide sequence (Supplementary Figure 1). Among these, clones with higher fluorescence intensity than previously developed translation enhancing SKIK peptide were summarized in Figure 2. While no peptide sample without any inserted peptide showed low intensity value 16, that of SKIK was 86. At the screening step, IFRC had the highest intensity followed by FSYD, VSVD, ILDW, ISMD, and SAAD. Sequence logos were generated for both the positive and negative sequences (Figure 2B). A comparison of the two logos revealed that the negative clones showed a relatively uniform distribution of amino acids at all positions. In contrast, the positive clones displayed a distinct higher frequency of D at the fourth position, suggesting its potential contribution in enhancing translation.

**Figure 1.**
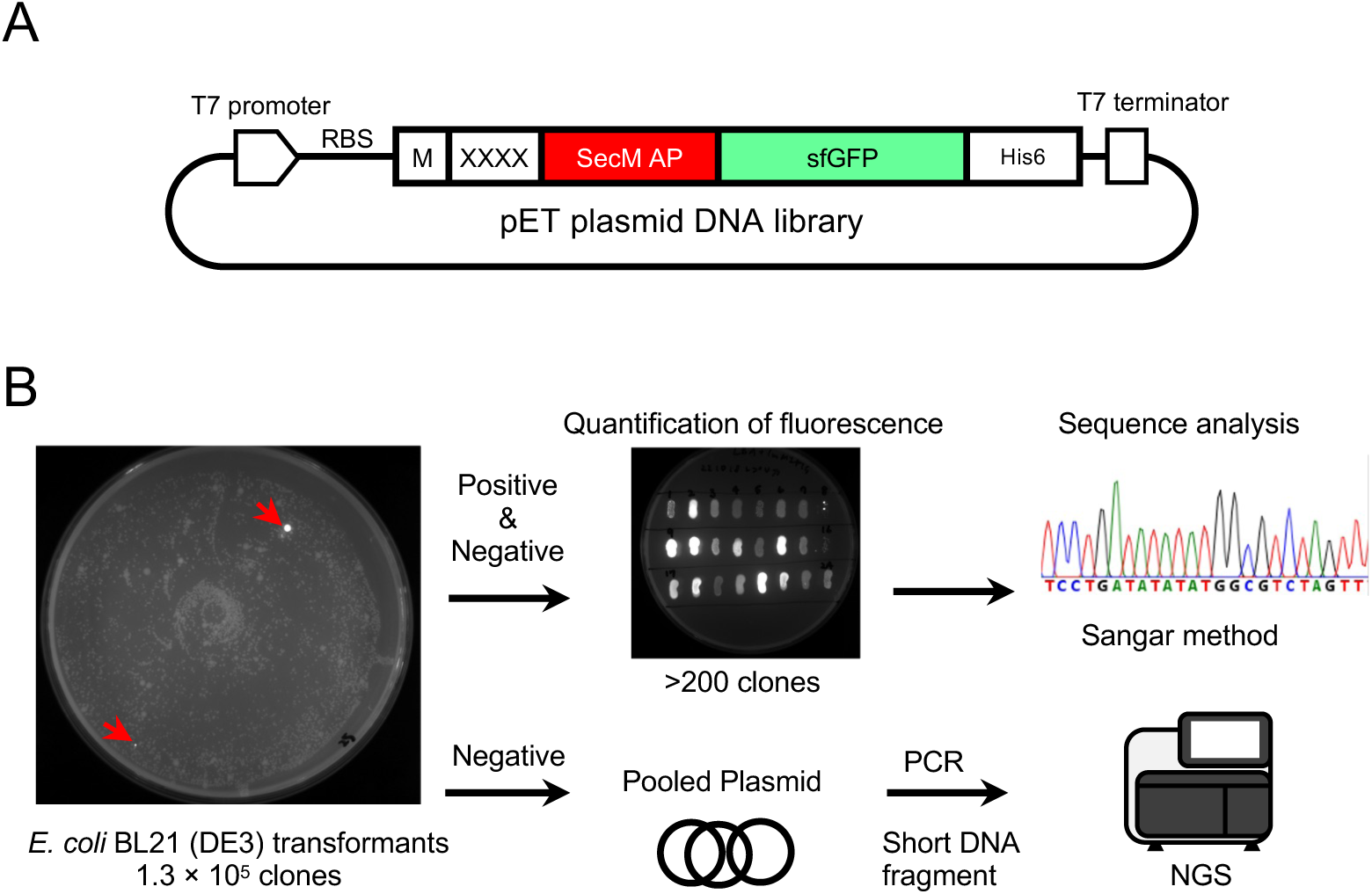
Screening scheme of translation enhancing peptides in *E. coli*. A. Plasmid library constructed in this study. RBS; ribosome binding site, XXXX; randomized peptide encoded by (NNK)_4_ sequence. B. Outline of screening procedure and sequence analysis. Typical culture plates are show herein. Red arrows indicate examples of positive clones showing fluorescence.

**Figure 2.**
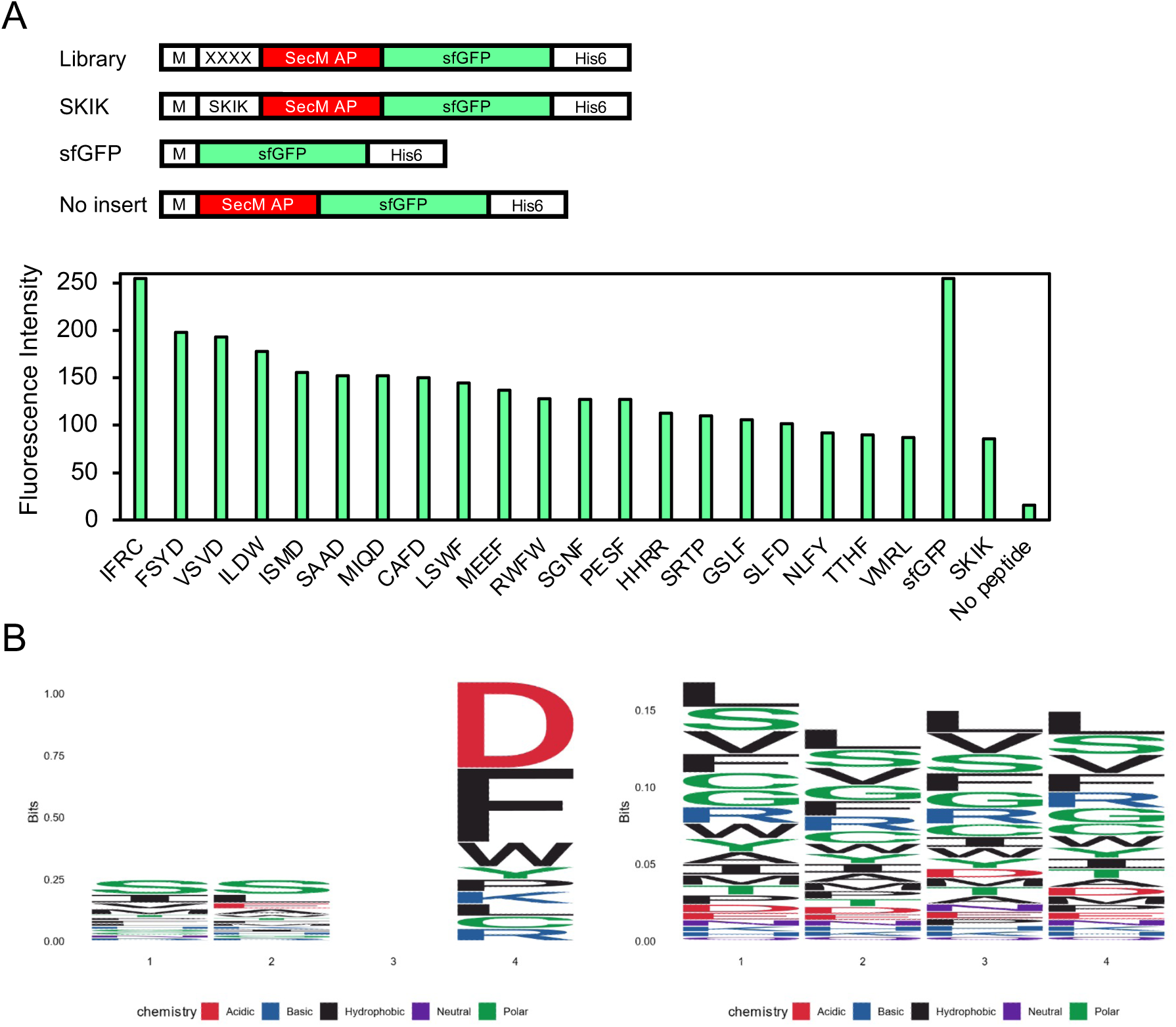
Analysis of the effect of fourth amino acid residues of FSYX. A. Relative fluorescence intensity of the picked *E. coli* positive clones on LB agar plates with 1 mM IPTG with stronger fluorescent than SKIK. Four amino acid sequence indicates the corresponding peptide XXXX in the library plasmid. DNA constructs of samples and controls are illustrated above. B. Logo plot analysis. Left and right panels indicate the result of the positive 20 clones exhibited stronger fluorescence than SKIK and negative clones analyzed by Nextseq 550, respectively.

### In vitro assay using PURE system

Based on the above analysis, clones with strong fluorescence—indicating a strong ability to counteract the ribosomal stalling by arrest of SecM AP—frequently had D as the fourth amino acid in the inserted tetrapeptide. Therefore, we experimentally investigated the importance of fourth amino acid on translational enhancement efficiency by substituting FSYD to FSYX (Supplementary Table 1). To analyze only the effect on translation, we used a reconstituted *E. coli* cell-free translation PURE system instead of a live cell expression system. The relative fluorescence intensity of each variant, normalized to 1 for the no peptide control (i.e., without tetrapeptide insertion), is shown in Figure 3A. As seen in the results, FSYD, FSYE, FYYN, and FSYQ showed high fluorescence intensities, followed by FSYK. We therefore plotted the physicochemical properties^34^ of the fourth amino acid against fluorescence intensity, and found that two parameters—side chain hydrophobicity and in/out-propensity—were inversely correlated with fluorescence intensity, with R² values exceeding 0.5 (Figure 3B).

**Figure 3.**
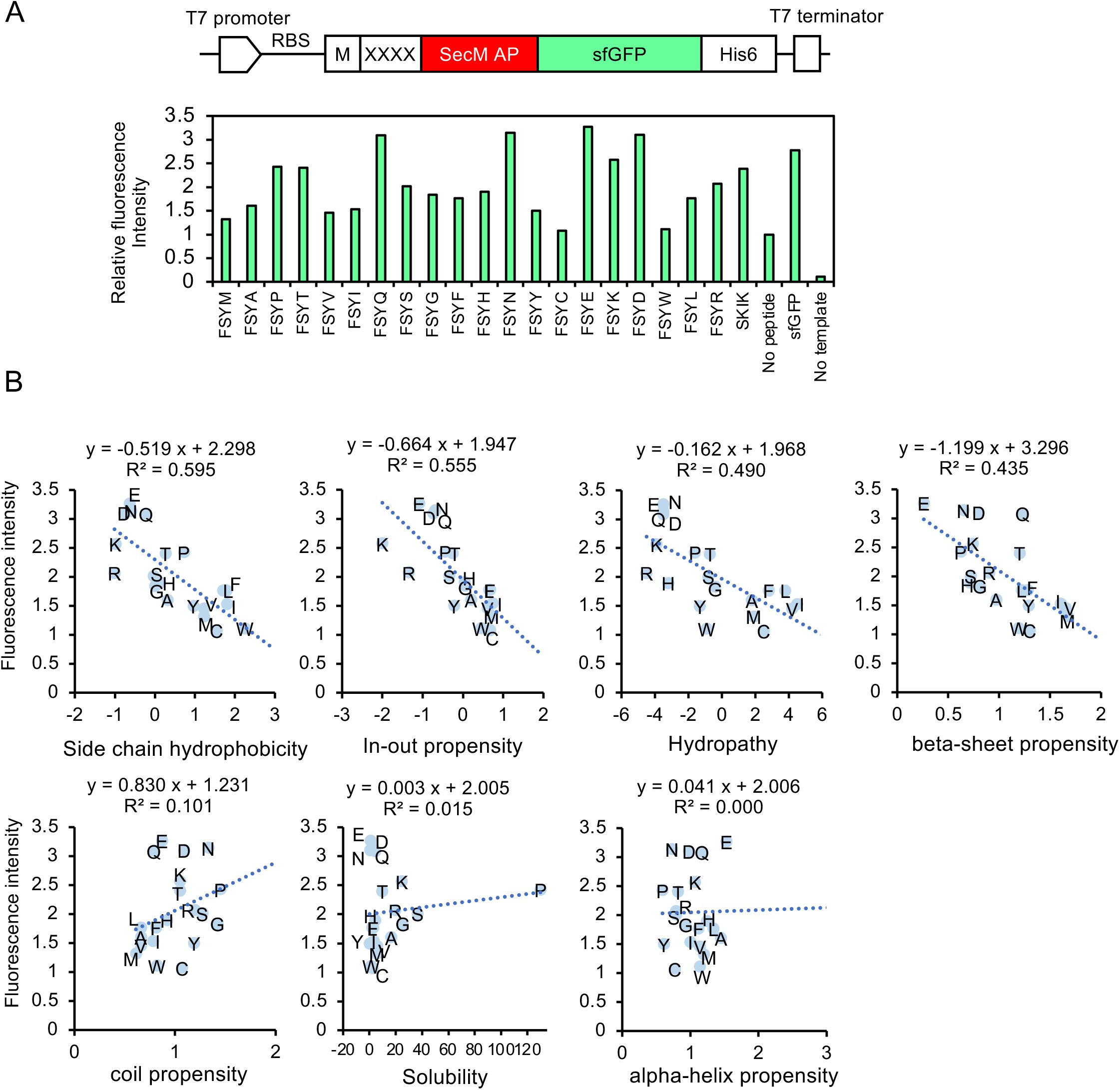
Influence of the fourth amino acid residue in FSYX peptide on alleviating ribosomal stalling by SecM AP. A. DNA construct used for CFPS and fluorescence analysis. The region XXXX is replaced with the peptide sequence shown here. No peptide means no XXXX insertion between M and SecM AP. sfGFP is positive control without XXXX-SecM AP. No template is negative control of CFPS without any DNA template. Fluorescence intensity of No peptide was regarded as 1. B. Various parameters of fourth amino acid amino of FSYX and relative fluorescence intensity. Seven parameters were cited from previous work by Nomoto et al^34^.

To determine whether the peptides identified through in vivo screening truly enhance translation, we evaluated all peptide sequences analyzed by Sanger sequencing using the PURE system, which is independent from cellular growth conditions and cellular background components. As a result, peptides such as VSVD, FSYD, SAAD, and ISMD— which showed high fluorescence intensity in vivo—also demonstrated strong translation enhancing effects in the in vitro system, with relative values of 1.3, 1.2, 1.2, and 1.0 respectively, when normalized to the translation level of SKIK as 1.0 (Figure 4). These results suggest that although the values from in vivo and in vitro systems do not completely correlate, many peptides with possible translation enhancing activity were successfully identified through this screening.

**Figure 4.**
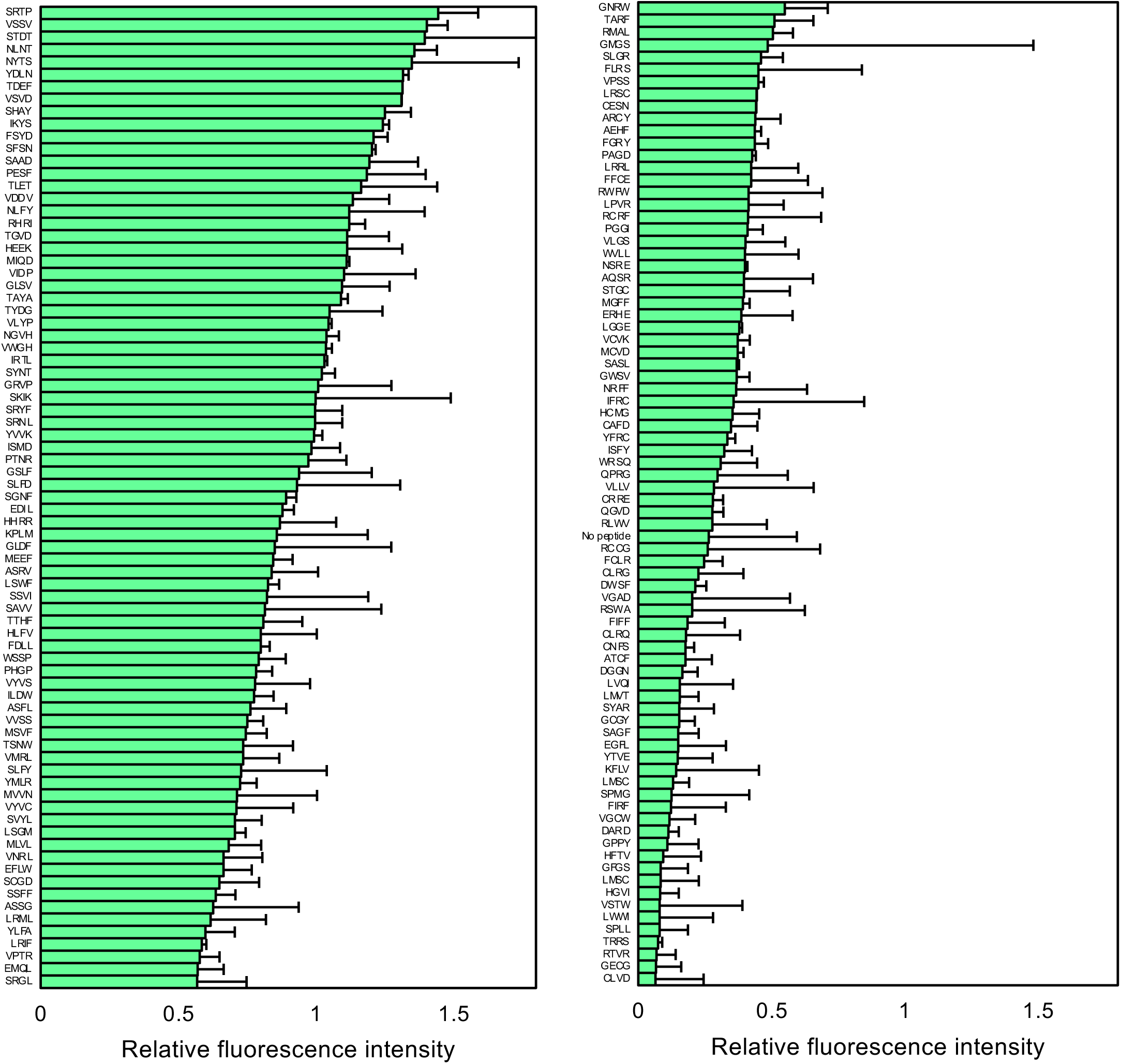
Quantitative fluorescence analysis of the in vitro translated products. The positive and randomly selected clones identified through in vivo screening were evacuated using CFPS. The relative fluorescence intensities, normalized by SKIK (regarded as 1), are presented. Error bars represent standard deviation of three independent experiments.

It is known that protein expression levels often differ between in vitro and in vivo systems due to differences in molecular environments, such as mRNA stability, folding efficiency, and the presence of regulatory or degradation machinery^35,36^.While certain 5′-UTR sequences enhanced protein expression in *E. coli* strains JM109 and BL21, these effects were not consistently replicated in a cell-free in vitro system^37^, suggesting it is challenging to identify factors that function universally across both in vitro and in vivo systems.

Trans-translation is reported as a quality control mechanism in bacteria that rescues stalled ribosomes on defective mRNAs^38–40^. It involves transfer-messenger RNA (tmRNA) and the protein SmpB, which together release the ribosome and tag the incomplete polypeptide for degradation. Importantly, the ribosome rescue system differs between in vivo and in vitro translation. In in vivo systems, such as in *E. coli* cells, trans-translation actively resolves ribosome stalling. However, in reconstituted in vitro systems like the PURE system, this rescue pathway is absent unless tmRNA and SmpB are supplemented. Therefore, translation stalling events may have different outcomes depending on the system used.

Strategies that involve modifying the N-terminus of the gene of interest or adding tetrapeptides to enhance the production of target proteins have been reported^21,41^. However, to the best of our knowledge, this is the first comprehensive screening that uses the apparent alleviation of translation stalling caused by APs as a selection criterion.

### First round bioinformatics analysis

To construct a TEP prediction model, we employed and compared two machine learning approaches: random forest ^42^ and XGBoost ^43^. Although XGBoost generally offers higher accuracy than random forest, it is prone to becoming trapped in local minima. In the initial training, a dataset comprising 158 sequences (157 newly designed peptides and SKIK) along with their in vitro translation-derived fluorescence intensities (Figure 4) was used. As direct learning from amino acid letters did not yield reliable predictions of brightness, we utilized established amino acid descriptors—Z-scale^44^, T-scale^45^, ST-scale^46^ VHSE-scale^47^, and EnsembleEnergy^48^—as explanatory variables. The performance of the first random forest model is summarized in Figure 5. The Pearson correlation coefficient and root mean square error (RMSE) between predicted and observed values for the overall model were 0.50 and 0.37, respectively (Figure 5A). Feature importance analysis revealed that position-independent descriptors such as Z-scale component 5 and T-scale component 3 were among the most influential features. EnsembleEnergy variables also ranked highly (Figure 5B). A sequence logo generated from the top 100 predicted sequences with high fluorescence indicated a strong preference for N as the first amino acid residue (Figure 5C). Additionally, a Sankey diagram illustrating patterns of adjacent amino acids revealed a dominant layout of peptide sequences (Figure 5D). The results of the XGBoost model are presented in Supplementary Figure 2. In this model, the Pearson correlation coefficient was 0.51, comparable to that of the random forest model.

**Figure 5.**
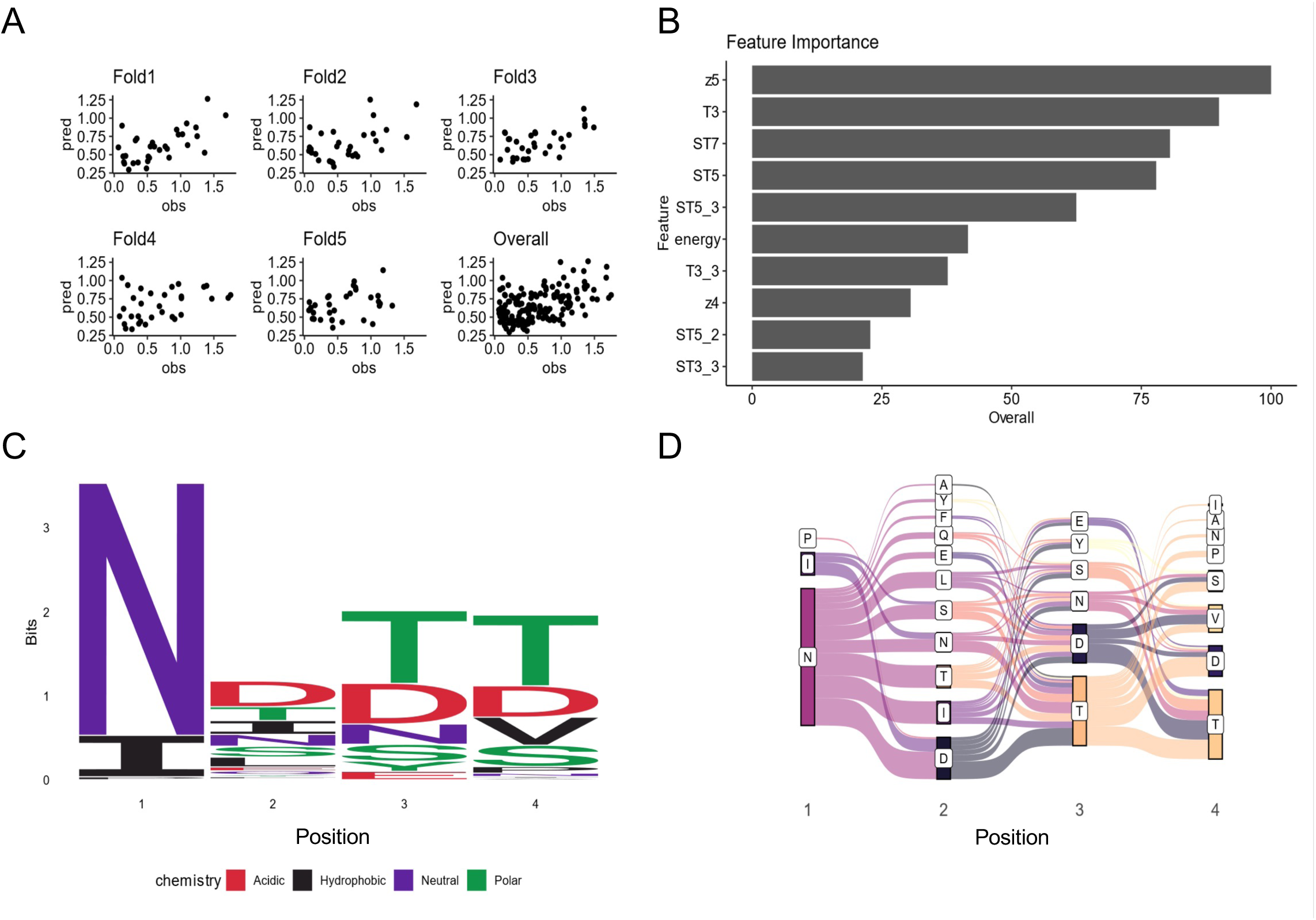
Analysis of in vitro data and first prediction of positive peptide sequences. Regression using 5-fold cross-validated random forest was performed, with the measured relative fluorescence intensity in vitro (normalized to a value of 1 for SKIK) as the response variable. The explanatory variables included z-scale, t-scale, st-scale, vhse-scale, and the ensemble energy of mRNA. A. Scatter plots of predicted values (pred) and observed values (obs) for each fold and across all folds. B. Top 10 most important explanatory variables in the trained random forest model. C. Sequence logo of the top 100 peptide sequences with the highest predicted brightness values. D. Sankey diagram of the top 100 peptides with highest predicted values.

### Establishing a loop to improve accuracy of prediction by incorporating new data

To evaluate the performance of the initial prediction model, 50 peptides with the highest predicted fluorescence intensities—out of 5,000 candidates generated by first-round random forest training—were experimentally assessed using *E. coli* reconstituted cell-free protein synthesis (CFPS) (Supplementary Table 2). The results demonstrated that many of these peptides exhibited translation enhancing activity comparable to that of SKIK (Figure 6A).

**Figure 6.**
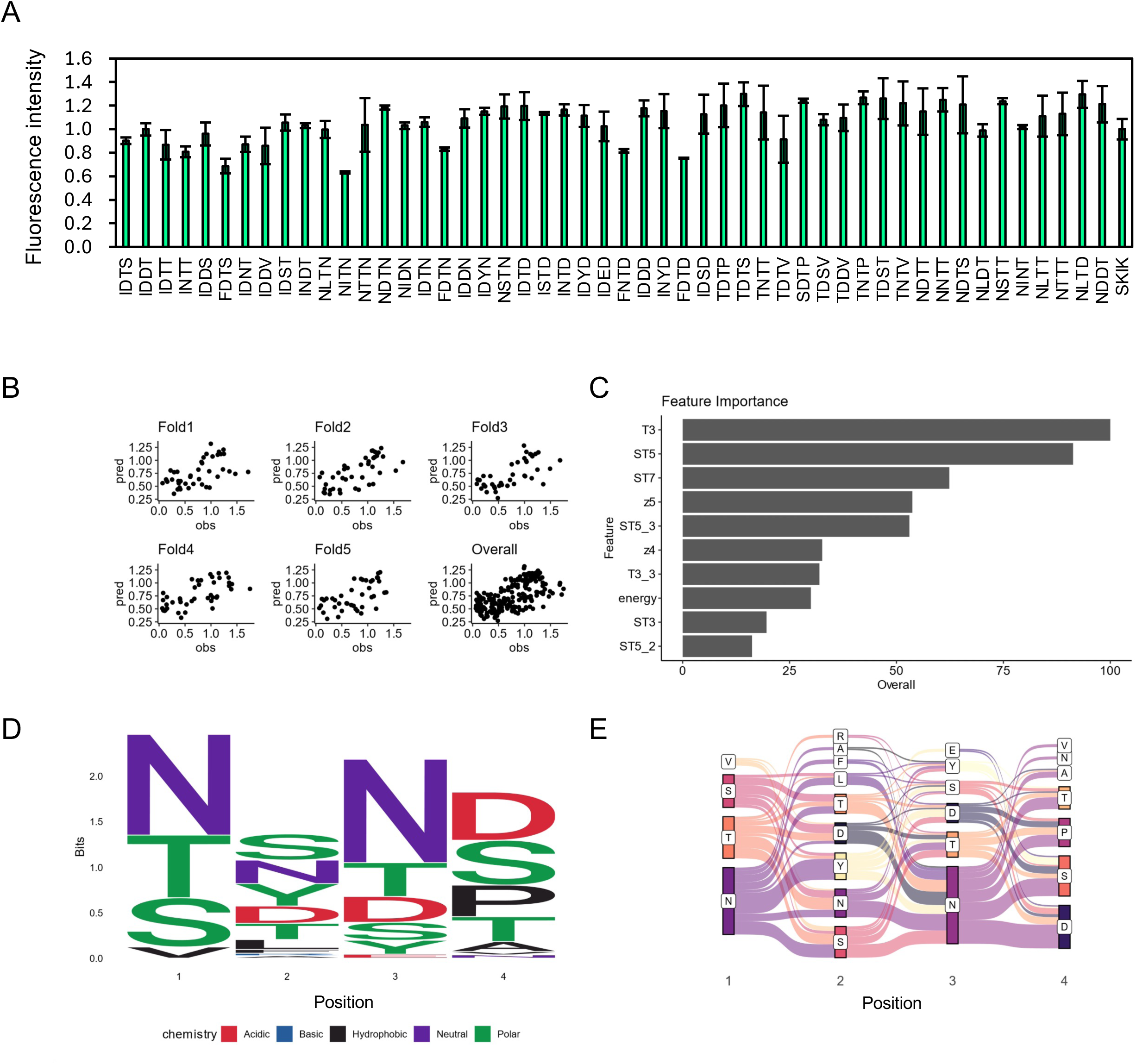
Analysis of in vitro data and second prediction by random forest model. A. Based on the first round of model training, 50 peptide sequences predicted to exhibit high fluorescence intensity were selected. Their translation-enhancing activities were evaluated by measuring sfGFP fluorescence in a CFPS system. Relative fluorescence intensities are shown, using SKIK as the reference standard with a value of 1. The corresponding peptide and DNA sequences are shown in Supplementary Table 2. B. Scatter plots of predicted values (pred) and observed values (obs) for each fold and all folds. C. Top 10 most important explanatory variables in the random forest model trained with the additional data from panel A. D. Sequence logo of the top 100 peptide sequences with the highest predicted fluorescence values. E. Sankey diagram of the top 100 peptide sequences with the highest predicted fluorescence values.

To improve the predictive accuracy, a second round of machine learning was conducted using the experimental data from the initially predicted 50 peptides, in addition to the original dataset of 158 sequences, as training data. After the second-round training, the overall correlation between predicted and experimentally measured fluorescence intensities across all cross-validation folds increased from 0.50 to 0.64 (Figure 5A, 6B). Similarly, the correlation coefficient for the XGBoost model improved from 0.51 to 0.63 (Supplementary Figure 3). Although random forest and XGBoost do not exactly coincide, they showed same trend.

The density maps in the Supplementary Figure 4, which show the predicted fluorescence intensities, indicate that the overall fluorescence is distributed mostly at lower intensities. This suggests that highly active translation enhancing peptides are relatively rare, which is consistent with the experimental results obtained from the screening.

Sequence logo analysis (Figure 6C) and Sankey diagram visualization (Figure 6D) of the top 100 predicted peptides revealed patterns consistent with those observed in the first round (Figure 5C). Notably, hydrophilic amino acids were favored across all positions, with D frequently enriched at positions 2 to 4.

To evaluate prediction accuracy after the second-round training, we selected the top 10 peptides with the highest predicted fluorescence from each of the random forest and XGBoost models. Additionally, 6 peptides highly ranked by both models and 15 randomly chosen sequences regardless of the predicted values were included for experimental validation (Supplementary Table 3). The measured fluorescence intensities are shown in Figure 7A, and their comparison with predicted values is shown in Figure 7B. A strong correlation (R = 0.83) was observed across both top-ranked and random sequences.

**Figure 7.**
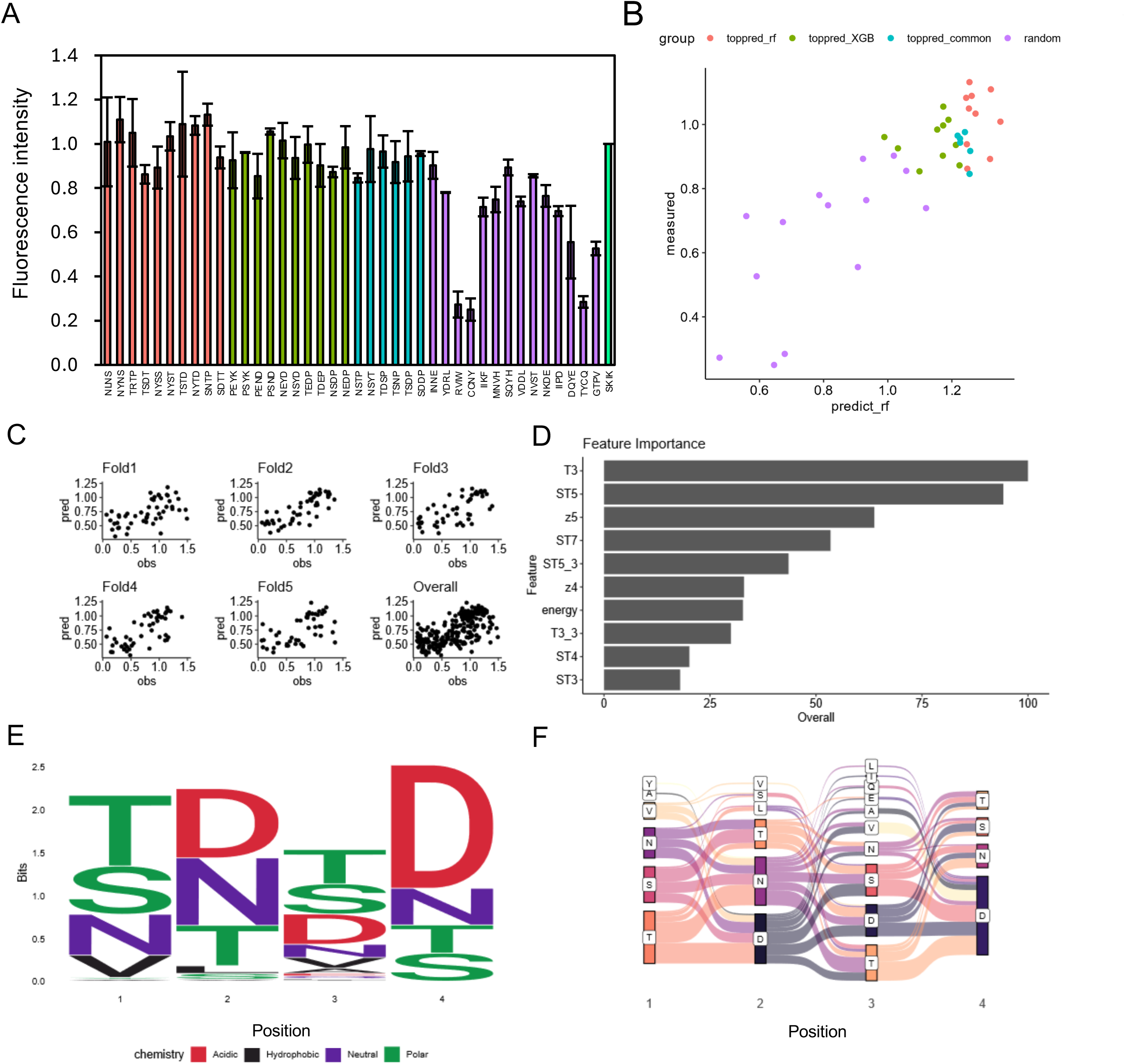
Analysis of various peptides predicted by second and third prediction by random forest model. A. Relative fluorescence intensities of sfGFP expressed in CFPS are shown for peptide sequences predicted by the second round of training: the top 10 sequences from random forest (red), the top 10 from XGBoost (green), the top 10 common to both models (cyan), and 14 sequences randomly selected regardless of predicted intensity (purple). Each peptide was fused to the N-terminus of SecM AP-sfGFP. The fluorescence intensity is shown relative to SKIK with a value of 1, with mean and standard deviation from triplicate experiments. B. The relationship between the predicted fluorescence intensities of output sequences from each of the two trained models and the experimentally measured values shown in panel A is presented. The color of each sample corresponds to the color of the bars in panel A. C. Scatter plots of predicted values (pred) and observed values (obs) for each fold and all folds. D. Top 10 most important explanatory variables in the random forest model trained with the additional data from panel A. E. Sequence logo of the top 100 peptide sequences with the highest predicted fluorescence values by random forest learning model. F. Sankey diagram of the top 100 peptide sequences with the highest predicted fluorescence values.

The third round of machine learning was performed using the second training dataset, supplemented with the experimentally measured values of the 50 peptides described above. The correlation coefficient improved to 0.66 in the random forest model (Figure 7C) and to 0.65 in the XGBoost model (Supplementary Figure 5), showing comparable performance. A summary of model accuracy across all rounds is provided in Supplementary Table 4.

The sequence logo of the top 100 predicted peptides revealed trends consistent with previous rounds: hydrophilic residues were favored at all positions, with D frequently enriched at positions 2–4, particularly at position 4 (Figure 7E). The Sankey diagrams also consistently highlighted preferred dipeptide motifs such as NN, SN, NS, ND, NT, and NP (Figures 5C, 6D, and 7F). The consistent patterns observed in the sequence logos and Sankey diagrams across all three rounds suggest that the models successfully identified common physicochemical features that define translation enhancing peptides.

### Feature analysis of TEPs predicted by trained models

Across all three rounds of machine learning, the top 10 most important features among the 157 amino acid descriptors remained largely consistent (Figure 5B, 6C, 7D). In the first round, key features associated with translation enhancing activity included z5, T3, and ST7. In the second and third rounds, T3, ST5, ST7, and z5 consistently ranked among the top contributors. T-scales summarize 67 topological descriptors related to amino acid connectivity into five principal components, explaining 91.1% of the variance. ST-scales expand on this by incorporating 827 3D structural features, compressed into eight components accounting for 71.5% of the variance. These scales, along with MS-WHIM, are known to exhibit similar behavior in capturing amino acid similarity^49^. Z-scales, derived from experimental data such as NMR and thin-layer chromatography, represent properties including lipophilicity (Z1), bulk (Z2), polarity/charge (Z3), and more complex characteristics like electronegativity and electrophilicity (Z4, Z5). Since all of these scales are derived through principal component analysis, direct interpretation of individual components is often not straightforward. However, they are widely used to reflect amino acid similarity and behavior in a compact, informative form.

Across three rounds of random forest modeling, the correlation between predicted and measured fluorescence values steadily improved, indicating enhanced predictive power as training data accumulated. Notably, certain features such as T3, ST5, and z5 consistently ranked among the most important variables throughout all rounds. This stability in feature selection supports the reliability and robustness of the model, even in the presence of descriptor redundancy.

In this study, machine learning enabled prediction of TEP candidates from a vast sequence space of 160,000 possibilities, using only a small experimentally measured dataset. Starting with a low-bias training set, iterative model updates with predicted high-performing sequences improved accuracy. Given the unknown mechanism of TEPs and limited data, interpretable models like random forest and XGBoost were well-suited. This strategy efficiently narrowed the search space and identified novel TEPs that could not be obtained through experimental screening alone.

## Methods

### Construction of the plasmid library

pET22b-SecM AP-sfGFP (constructed in our previous study^18^) was used as the PCR template. To insert (NNK) × 4 codons immediately after the initiation codon ATG, inverse PCR was performed with primer pair, 5’-GGAGATATACATATG<u>NNKNNKNNKNNK</u>TTCAGCACGCCCGTCTGGATAAG-3’ and 5’-CATATGTATATCTCCTTCTTAAAGTTAAAC-3’ by KOD One polymerase (Toyobo, Osaka, Japan). The underlined nucleotides correspond to the randomized four amino acids. The PCR product was treated with DpnI (Takara Bio, Kusatsu, Japan, 37°C for 30 min and 70°C for 10 min for inactivation) and purified with the spin column (Econospin, Ajinomoto Bio-Pharma, San Diego, CA). The purified linear vector DNA was then self-assembled by Gibson assembly (New England Biolabs, Ipswich, MA). High performance *E. coli* HST08 competent cells (Takara Bio) were transformed with the product and colonies grown on LB plates containing 100 mg/L ampicillin were pooled with TE buffer (10 mM Tris-HCl, 1 mM EDTA, pH8.0) followed by the plasmid extraction using the commercial plasmid purification kit (Plasmid DNA Extraction Midi kit, Favorgen biotech corp, Ping Tung, Taiwan).

### Screening of *E. coli* clones

The above obtained plasmid library “pET22b-(NNK)_4_-SecM AP-sfGFP” was introduced to *E. coli* BL21 (DE3) strain for protein expression and spread on LB agar plates containing 100 mg/L ampicillin and 1 μM isopropyl β-D-thiogalactopyranoside (IPTG). The chemical competent cells were prepared with the commercially available kit (Mix & Go, Zymoresearch, Irvine, CA). The colonies grown on the LB plate at 37°C for 16 h were analyzed by the fluorescent imaging apparatus MultiImager II with the appended software MISIS II (BioTools Co., Ltd., Takasaki, Japan, filter set ex 485 nm, em 590/60nm band-pass, gain setting 13 dB, exposure 16 mS). The picked single colonies were suspended in 50 µL of sterile water and further spotted onto fresh culture plates as same as above and incubated another 16 h at 37°C. The brightness of the fluorescence from the growing colonies was analyzed by ImageJ software.

### Sequence analysis

Sequence analysis of all the positive clones with fluorescence and some negative clones exhibiting low fluorescent was performed by Sanger method using colony-directed PCR products amplified with the primer F1 (ATCTCGATCCCGCGAAATTAATACG) and R1 (TCCGGATATAGTTCCTCCTTTCAG) which anneal to upstream of T7 promoter and downstream of T7 terminator, respectively. Remaining *E. coli* clones which did not exhibit significant fluorescence were regarded as negative ones. They were suspended with TE buffer and pooled from the agar plates and their plasmids were extracted by plasmid extraction midi kit as described above. The DNA fragment (161 bp) containing randomized region was amplified with the primer pair AAGAAGGAGATATACATATG and ATTAACATCACCATCCAGTTC from the extracted plasmid as the template. The purified DNA fragment was then analyzed with Nextseq 550 by single-end read mode (81 bp) to fully cover the randomized region. NEBNext Ultra II DNA Library Prep Kit for Illumina and NextSeq 500/550 High Output Kit v2 (75 cycles) were used for sample preparation. The obtained data was analyzed by Seqkit^50^ and python program to get peptide sequence list of the negative clones.

### In vitro protein expression

To confirm the effect of the obtained peptides on translation, cell-free protein synthesis (CFPS) was conducted using PUREfrex2.1 (GeneFrontier, Kashiwa, Japan) with the DNA fragments amplified using Gflex DNA polymerase (Takara) with F1 and R1 primers from single colonies. CFPS condition was as follows; DNA template; 1 μL, Solution I; 2 μL, Solution II; 0.125 μL, Solution III (ribosome); 0.5 μL, and DEPC treated RNase free water (Nacalai Tesque, Kyoto, Japan); 1.375 μL. Reaction was performed at 37°C for 90 min with triplicate. Each CFPS reaction solution was diluted 25-fold with water and 50 µL was dispensed into the wells of Black Microplate Flat Bottom 96Well (Stem, Hino, Japan), and the fluorescence intensity of sfGFP was measured by a microplate reader (Infinite 200 PRO, TECAN, ZH, Switzerland) at excitation wavelength 485 nm (bandwidth 9 nm)/emission wavelength of 535 nm (bandwidth 20 nm).

### Construction of the mutants

The peptide FSYX (X means 20 amino acid) and another peptide candidates obtained by machine leaning prediction tool were introduced by PCR using KOD One (Toyobo) with the corresponding primer pairs as shown in Supplementary Table 1. *E. coli* HST08 competent cells were transformed with the *Dpn*I treated amplified PCR products. The plasmids with correct sequences were purified and used for further experiment.

### Data mining from in vivo data

A computational analysis was performed to find the characteristics of the TEPs obtained by in vivo screening. The peptide sequences with identical amino acid compositions and their corresponding fluorescence intensity values were averaged and combined into a single data point. To investigate sequence features associated with translation enhancing activity, we first analyzed the sequence composition of both positive and negative clones. Sequences were classified as positive if their fluorescence intensity exceeded that of the SKIK control. Sequence logos were then generated using the ggseqlogo package in R to visualize amino acid preferences at each position.

### Prediction of TEPs with machine learning

We next utilized in vitro experimental data to train models for predicting novel TEP sequences with high fluorescence. All fluorescence values used for training were normalized relative to the SKIK sequence. The input features included four amino acid descriptor sets—Z-scale, T-scale, ST-scale, and VHSE-scale—which numerically represent physicochemical and geometric properties of amino acids. For each descriptor, values were calculated for each residue position and the overall sequence average. In addition, to account for codon-level effects on translation, the mRNA free energy of the first 11 codons was calculated using the EnsembleEnergy function from RNAstructure, as described in a previous study^51^.

The Z-scale, T-scale, ST-scale, and VHSE-scale descriptor sets contain 5, 5, 8, and 8 features, respectively. Each feature set was applied to six aspects of the peptide sequence: the five individual residue positions (including the N-terminal methionine) and the overall sequence average. This resulted in a total of 156 amino acid–based features. Additionally, we included the EnsembleEnergy value of the first 11 codons as an mRNA-level descriptor, bringing the total number of features used in model training to 157. To enhance clarity given the large number of features, simplified abbreviations are used throughout this study. For example, the average value of a descriptor across the entire sequence is labeled as “z1,” following the original notation. In contrast, values corresponding to specific positions within a sequence include a positional suffix, such as “z1_3,” where “3” indicates the position in the peptide sequence. Position numbering begins at 0, with “0” referring to the initial M residue. The descriptor “EnsembleEnergy” is abbreviated simply as “energy.”

A 5-fold cross validation was performed to ensure robustness of the analysis. Random forest and XGBoost models were trained by the caret package of the R language. The ntree parameter in the random forest was set to 1,000 to ensure sufficient convergence of the calculation, and the mtry parameter was determined by grid search techniques. The optimal model was evaluated using predicted and measured RMSEs. Similarly, in XGBoost, the optimal value of eta, max_depth, gamma, colsample_bytree, min_child_weight, subsample, nrounds parameter were determined by grid search. In each round, the model with the lowest RMSE was selected using grid search. However, since RMSE values can be influenced by differences in the range or distribution of the training data, Pearson’s correlation coefficient was used to evaluate and compare model performance across rounds. Unlike RMSE, the correlation coefficient is scale-independent and reflects the strength of the linear relationship between predicted and observed values, making it more suitable for comparisons across datasets of varying sizes. All the optimal values for the hyperparameters are shown in Supplementary Table 5.

After training, fluorescence intensity for all possible tetrapeptide sequences (approximately 160,000 combinations) was predicted and TEP candidate sequences were selected. For experimentally verifying these predicted peptides, optimal codons were assigned based on the *E. coli* codon adaptation index^52^. To maximize sequence diversity, candidates were selected not only based on top predicted scores but also by clustering to ensure a broad representation of sequence types. In our experimental process, an iterative training approach was employed, consisting of three rounds of machine learning and experimental validation. In the first round, 50 high-scoring candidate peptides were selected based on the trained model and evaluated by in vitro translation. The measured data were then incorporated into the training dataset for the second round. An additional 40 peptides were experimentally tested and included in the third round of training. This iterative process was designed to improve both the accuracy of the predictive models and the efficiency of candidate identification. Model performance was assessed in each round to monitor improvements in predictive accuracy.

## Supporting information

The supporting information including all supplementary tables and figures is available free of charge.

## Author contributions

T.O.K. and G.Y equally contributed this work.

T.O.K. and H.N designed the study.

G.Y., M.H., and C.M. performed bioinformatic analysis and developed prediction tool.

T.O.K performed experiments.

T.O.K., G.Y., M.H., and C.M. analyzed the data.

T.O.K., G.Y., M.H., and C.M. wrote the manuscript.

T.O.K. supervised the project.

## Acknowledgment

This research was supported by AIST-Nagoya University alliance project, Japan Science and Technology agency FOREST Program (grant No. JPMJFR2204), GteXProgram Japan Grant Number JPMJGX23B6 and JPMJGX23B4. A part of this work was also supported by Japan Society for the Promotion of Science KAKENHI 23K04989. PURE*frex* 2.1 and supplemental components used in this study were kindly provided by GeneFrontier Corporation. We also acknowledge to Mrs. Harumi Masuda for her technical support.

## Abbreviations

Aps: arrest peptides
CFPS: cell-free protein synthesis
TEP: translation-enhancing peptide
POI: protein of interest
sfGFP: superfolder green fluorescent protein
IPTG: isopropyl β-D-thiogalactopyranoside

## Notes

### Competing Interest Statement

The authors have declared no competing interest.

