## supporting information for "Screening and machine-learning assisted prediction of translation-enhancing peptides reducing ribosomal stalling in *Escherichia coli*"

Supplemental Table 1. DNA sequences corresponding to FSYX

| Peptide | DNA | Primers used |
|---|---|---|
| FSYD | TTTAGTTATGAT | n/a because this sequence was obtained in the screening |
| FSYW | TTTAGTTATTGG | catATGTTTAGTTATTGGTTCAGCACGCCCGTCTGGATAAGC<br>TGCTGAACCAATAACTAAACATATGTATATCTCCTTCTTAAAG |
| FSYL | TTTAGTTATTTA | catATGTTTAGTTATTTATTCAGCACGCCCGTCTGGATAAGC<br>TGCTGAATAAATAACTAAACATATGTATATCTCCTTCTTAAAG |
| FSYR | TTTAGTTATAGA | catATGTTTAGTTATAGATTCAGCACGCCCGTCTGGATAAGC<br>TGCTGAATCTATAACTAAACATATGTATATCTCCTTCTTAAAG |
| FSYY | TTTAGTTATTAT | TAGTTATTATTTCAGCACGCCCGTCTGGATAAGC<br>TGCTGAAATAATAACTAAACATATGTATATCTCCTTC |
| FSYC | TTTAGTTATTGT | TAGTTATTGTTTCAGCACGCCCGTCTGGATAAGC<br>TGCTGAAACAATAACTAAACATATGTATATCTCCTTC |
| FSYE | TTTAGTTATGAA | TAGTTATGAATTCAGCACGCCCGTCTGGATAAGC<br>TGCTGAATTCATAACTAAACATATGTATATCTCCTTC |
| FSYK | TTTAGTTATAAA | TAGTTATAAATTCAGCACGCCCGTCTGGATAAGC<br>TGCTGAATTTATAACTAAACATATGTATATCTCCTTC |
| FSYQ | TTTAGTTATCAA | TAGTTATCAATTCAGCACGCCCGTCTGGATAAGC<br>TGCTGAATTGATAACTAAACATATGTATATCTCCTTC |
| FSYS | TTTAGTTATTCT | TAGTTATTCTTTCAGCACGCCCGTCTGGATAAGC<br>TGCTGAAAGAATAACTAAACATATGTATATCTCCTTC |
| FSYG | TTTAGTTATGGT | TAGTTATGGTTTCAGCACGCCCGTCTGGATAAGC<br>TGCTGAAACCATAACTAAACATATGTATATCTCCTTC |
| FSYF | TTTAGTTATTTT | TAGTTATTTTTTCAGCACGCCCGTCTGGATAAGC<br>TGCTGAAAAAATAACTAAACATATGTATATCTCCTTC |
| FSYH | TTTAGTTATCAT | TAGTTATCATTTCAGCACGCCCGTCTGGATAAGC<br>TGCTGAAATGATAACTAAACATATGTATATCTCCTTC |
| FSYN | TTTAGTTATAAT | TAGTTATAATTTCAGCACGCCCGTCTGGATAAGC<br>TGCTGAAATTATAACTAAACATATGTATATCTCCTTC |
| FSYM | TTTAGTTATATG | TAGTTATATGTTCAGCACGCCCGTCTGGATAAGC<br>TGCTGAACATATAACTAAACATATGTATATCTCCTTC |
| FSYA | TTTAGTTATGCT | TAGTTATGCTTTCAGCACGCCCGTCTGGATAAGC<br>TGCTGAAAGCATAACTAAACATATGTATATCTCCTTC |
| FSYP | TTTAGTTATCCA | TAGTTATCCATTCAGCACGCCCGTCTGGATAAGC |

| | | |
|---|---|---|
| | | TGCTGAATGGATAACTAAACATATGTATATCTCCTTC |
| FSYT | TTTAGTTATACA | TAGTTATACATTCAGCACGCCCGTCTGGATAAGC<br>TGCTGAATGTATAACTAAACATATGTATATCTCCTTC |
| FSYV | TTTAGTTATGTT | TAGTTATGTTTTCAGCACGCCCGTCTGGATAAGC<br>TGCTGAAAACATAACTAAACATATGTATATCTCCTTC |
| FSYI | TTTAGTTATATA | TAGTTATATATTCAGCACGCCCGTCTGGATAAGC<br>TGCTGAATATATAACTAAACATATGTATATCTCCTTC |

Supplementary Table 2. Predicted and measured peptides on the first-round training

| Peptide | DNA |
|---|---|
| IDTS | ATCGACACCTCT |
| IDDT | ATCGACGACACC |
| IDTT | ATCGACACCACC |
| INTT | ATCAACACCACC |
| IDDS | ATCGACGACTCT |
| FDTS | TTCGACACCTCT |
| IDNT | ATCGACAACACC |
| IDDV | ATCGACGACGTT |
| IDST | ATCGACTCTACC |
| INDT | ATCAACGACACC |
| NLTN | AACCTGACCAAC |
| NITN | AACATCACCAAC |
| NTTN | AACACCACCAAC |
| NDTN | AACGACACCAAC |
| NIDN | AACATCGACAAC |
| IDTN | ATCGACACCAAC |
| FDTN | TTCGACACCAAC |
| IDDN | ATCGACGACAAC |
| IDYN | ATCGACTACAAC |
| NSTN | AACTCTACCAAC |
| IDTD | ATCGACACCGAC |
| ISTD | ATCTCTACCGAC |
| INTD | ATCAACACCGAC |
| IDYD | ATCGACTACGAC |
| IDED | ATCGACGAAGAC |
| FNTD | TTCAACACCGAC |
| IDDD | ATCGACGACGAC |
| INYD | ATCAACTACGAC |
| FDTD | TTCGACACCGAC |
| IDSD | ATCGACTCTGAC |
| TDTP | ACCGACACCCCG |
| TDTS | ACCGACACCTCT |
| TNTT | ACCAACACCACC |

| | |
|---|---|
| TDTV | ACCGACACCGTT |
| SDTP | TCTGACACCCCG |
| TDSV | ACCGACTCTGTT |
| TDDV | ACCGACGACGTT |
| TNTP | ACCAACACCCCG |
| TDST | ACCGACTCTACC |
| TNTV | ACCAACACCGTT |
| NDTT | AACGACACCACC |
| NNTT | AACAACACCACC |
| NDTS | AACGACACCTCT |
| NLDT | AACCTGGACACC |
| NSTT | AACTCTACCACC |
| NINT | AACATCAACACC |
| NLTT | AACCTGACCACC |
| NTTT | AACACCACCACC |
| NLTD | AACCTGACCGAC |
| NDDT | AACGACGACACC |

Supplementary Table 3. Predicted and measured peptides on the second-round training

| Peptide | DNA | Prediction |
|---|---|---|
| NLNS | AACCTGAACTCT | Highest by random forest |
| NYNS | AACTACAACTCT | |
| TRTP | ACCCGTACCCCG | |
| TSDT | ACCTCTGACACC | |
| NYSS | AACTACTCTTCT | |
| NYST | AACTACTCTACC | |
| TSTD | ACCTCTACCGAC | |
| NYTD | AACTACACCGAC | |
| SNTP | TCTAACACCCCG | |
| SDTT | TCTGACACCACC | |
| PEYK | CCGGAATACAAA | Highest by random XGBoost |
| PSYK | CCGTCTTACAAA | |
| PEND | CCGGAAAACGAC | |
| PSND | CCGTCTAACGAC | |
| NEYD | AACGAATACGAC | |
| NSYD | AACTCTTACGAC | |
| TEDP | ACCGAAGACCCG | |
| TDEP | ACCGACGAACCG | |
| NSDP | AACTCTGACCCG | |
| NEDP | AACGAAGACCCG | |
| NSTP | AACTCTACCCCG | Highly ranked by both random forest and XGBoost |
| NSYT | AACTCTTACACC | |
| TDSP | ACCGACTCTCCG | |
| TSNP | ACCTCTAACCCG | |
| TSDP | ACCTCTGACCCG | |
| SDDP | TCTGACGACCCG | |
| INNE | ATCAACAACGAA | Randomly chosen sequences regardless of the predicted values |
| YDRL | TACGACCGTCTG | |
| RVIW | CGTGTTATCTGG | |
| CCNY | TGCTGCAACTAC | |
| IIKF | ATCATCAAATTC | |
| MNVH | ATGAACGTTCAC | |
| SQYH | TCTCAGTACCAC | |

| | |
|---|---|
| VDDL | GTTGACGACCTG |
| NVST | AACGTTTCTACC |
| NKDE | AACAAAGACGAA |
| IIPD | ATCATCCCGGAC |
| MDQYE | GACCAGTACGAA |
| TYCQ | ACCTACTGCCAG |
| GTPV | GGTACCCCGGTT |

Supplementary Table 4. Summary of evaluation metrics for each random forest and XGBoost model.

| Round | RF_cor | RF_RMSE | XGB_cor | XGB_RMSE |
|---|---|---|---|---|
| First | 0.50 | 0.37 | 0.51 | 0.37 |
| Second | 0.64 | 0.32 | 0.63 | 0.33 |
| Third | 0.66 | 0.28 | 0.65 | 0.28 |

Cor represents the Pearson's correlation coefficient, and RMSE denotes the root mean square error between measured and predicted values. RF and XGB mean random forest and XGBoost, respectively.

Supplementary Table 5. Summary of hyperparameter-tuning results from grid search for each model

| Round | RF | XGB | | | | | | |
|---|---|---|---|---|---|---|---|---|
| | mtry | nrounds | max_depth | eta | gamma | colsample_bytree | min_child_weight | subsample |
| First | 75 | 125 | 3 | 0.025 | 0 | 0.7 | 1 | 1 |
| Second | 60 | 200 | 2 | 0.025 | 0 | 1 | 1 | 1 |
| Third | 40 | 125 | 5 | 0.025 | 0 | 0.4 | 1 | 1 |

To avoid overcomplicating the XGBoost optimization, the parameters gamma, min_child_weight, and subsample were fixed, as they primarily influence learning rate and model generalization.

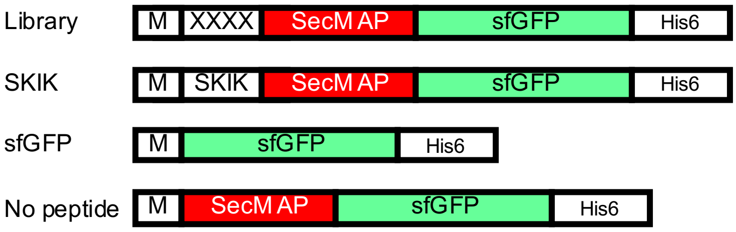


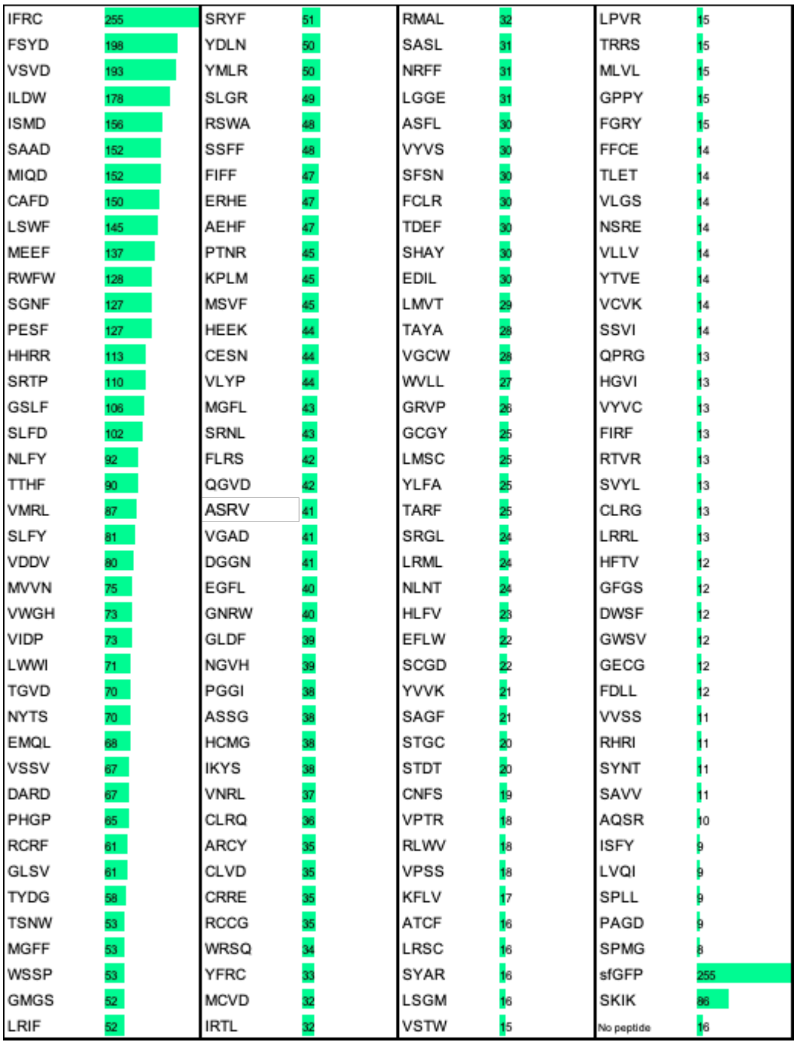


**Supplementary Figure 1. Analysis of the effect of four amino acid residues in *E. coli***

Relative fluorescence intensity of the picked *E. coli* clones on LB agar plates with 1 mM IPTG. Positive clones and randomized selected negative ones were analyzed. Four amino acid sequence indicates the corresponding peptide encoded by $(NNK)_4$ in the library plasmid. DNA constructs of samples and controls are illustrated above.

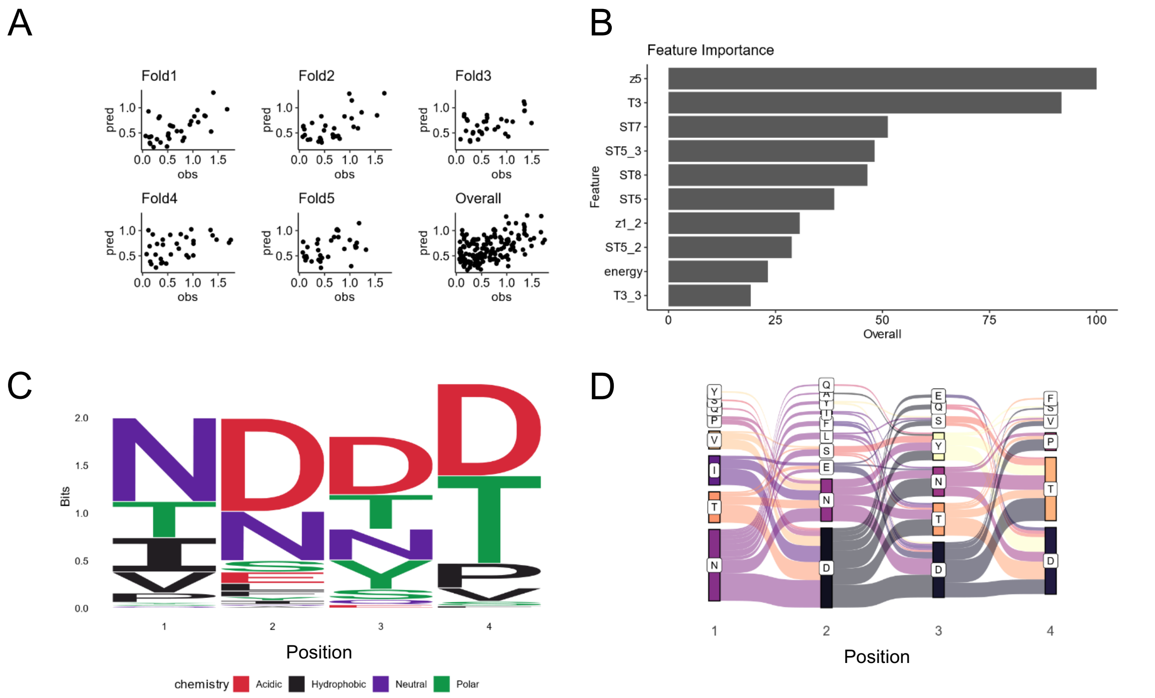


**Supplementary Figure 2. Analysis of in vitro data and first prediction of positive peptide sequences by XGBoost model**

Regression using 5-fold cross-validated XGBoost was performed, with the measured relative fluorescence intensity in vitro (normalized to a value of 1 for SKIK) as the response variable.

The explanatory variables included z-scale, t-scale, st-scale, vhse-scale, and the ensemble energy of mRNA.

A. Scatter plots of predicted values (pred) and observed values (obs) for each fold and across all folds.

B. Top 10 most important explanatory variables in the trained random forest model

C. Sequence logo of the top 100 peptide sequences with the highest predicted brightness values

D. Sankey diagram of the top 100 peptides with highest predicted values

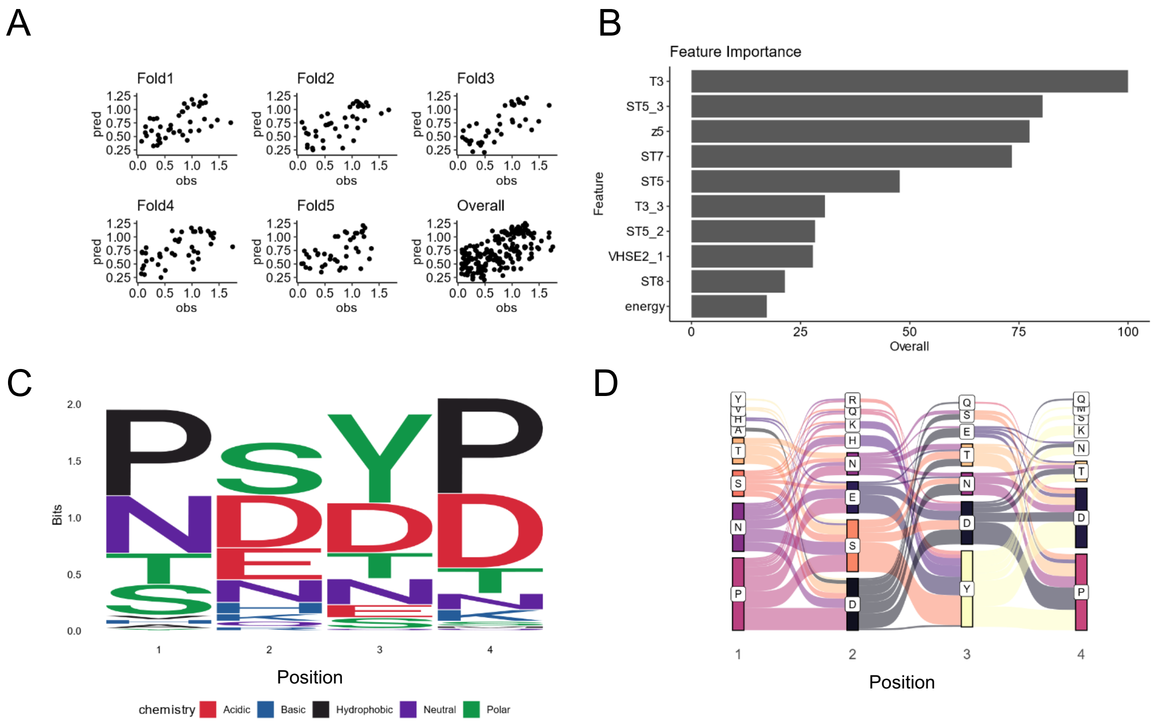


**Supplementary Figure 3. Analysis of in vitro data and second prediction by XGBoost model**

A. Scatter plots of predicted values (pred) and observed values (obs) for each fold and all folds.

B. Top 10 most important explanatory variables in the XGBoost model trained with the additional data from panel A.

C. Sequence logo of the top 100 peptide sequences with the highest predicted fluorescence values.

D. Sankey diagram of the top 100 peptide sequences with the highest predicted fluorescence values.

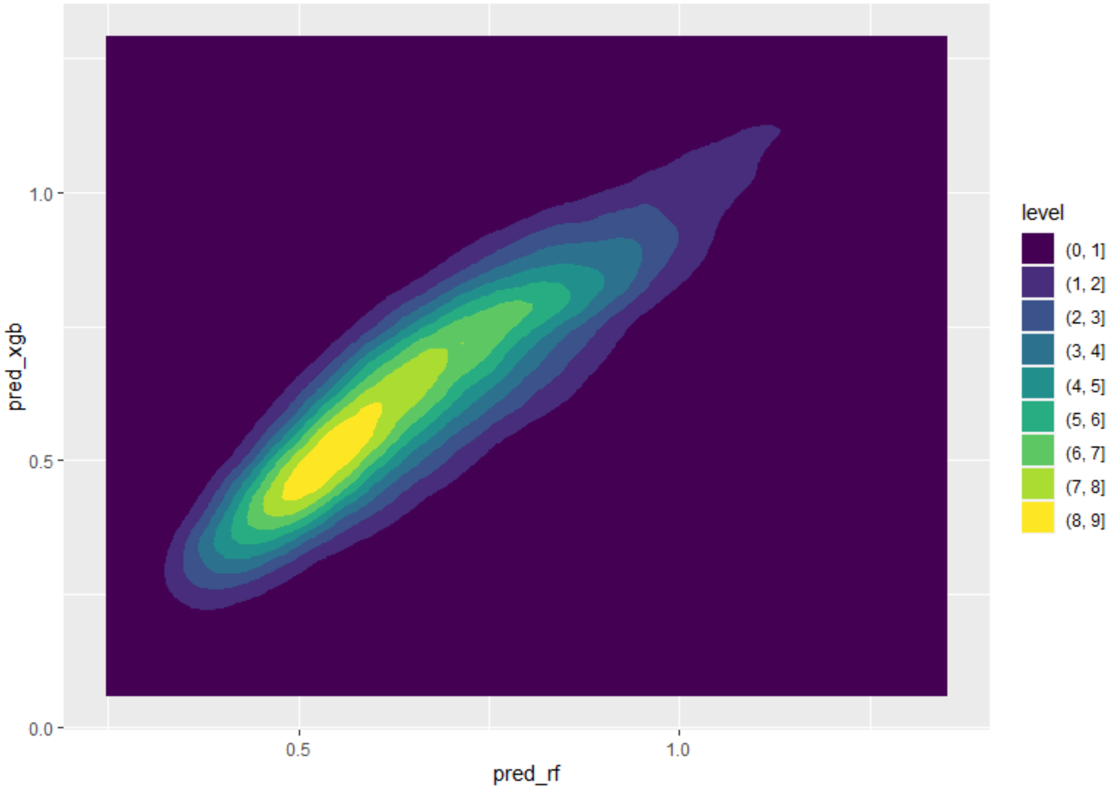


**Supplementary Figure 4. Contour plot comparing the predictions of both random forest and XGBoost from the second prediction**

Predicted values for all 160,000 possible tetrapeptides excluding measured ones are shown as a 2D kernel density plot, normalized to SKIK = 1. Color indicates density levels. Distributions predicted by random forest (Pred_rf) and XGBoost (Pred_xgb) are shown.

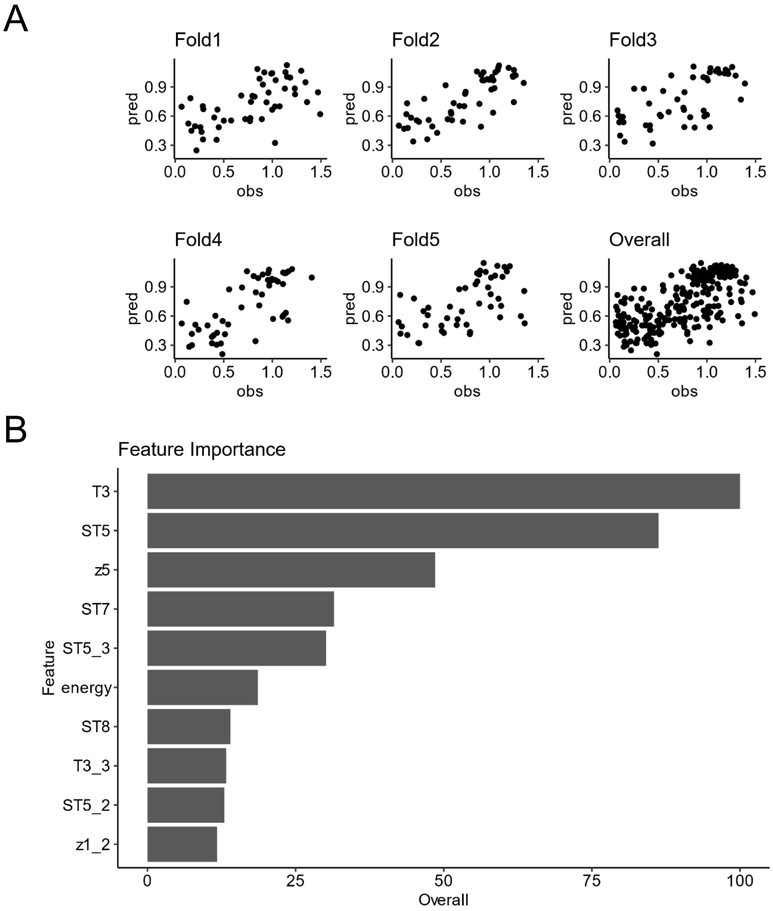


**Supplementary Figure 5. Analysis of in vitro data and third prediction by XGBoost model**

A. Scatter plots of predicted values (pred) and observed values (obs) for each fold and all folds.

B. Top 10 most important features identified in the XGBoost model trained with the expanded dataset.
